# Dental Age Assessment of White-tailed Deer via Computer Vision and Deep Learning

**DOI:** 10.1101/2025.07.25.666691

**Authors:** Aaron J. Pung

## Abstract

Accurate age estimation of white-tailed deer remains challenging for wildlife management, with existing computer vision methods limited to trail camera imagery and manual dental analysis requiring specialized expertise. This research presents the first computer vision approach to dental age assessment of white-tailed deer, eliminating the need for manual tooth replacement and wear evaluation. A dataset of 243 jawbone images is used to develop a multi-fold Convolutional Neural Network ensemble, achieving 90.7% ± 2.6% accuracy through 5-fold cross-validation. Attention map analysis confirms the model locates and focuses exclusively on dental features rather than imaging artifacts, successfully identifying the same wear patterns and eruption sequences that wildlife biologists rely upon in manual assessments. This implementation substantially outperforms computer vision models based on trail camera imagery and meets the accuracy standards required for scientific research and population management. The automated approach developed in this study enables rapid, objective age determination from dental specimens, dramatically reducing analysis time while offering immediate practical value for wildlife agencies, research institutions, and hunting programs requiring precise age data from harvested deer.

## 1 Introduction

### 1.1 Aging On The Hoof

Tracking and monitoring white-tailed deer populations is critical to the success of wildlife management. In part, this is due to the increasing importance of age-related data since deer age affects body growth, doe fertility, antler quality, and sex ratios of offspring.

State-of-the-art trail cameras enable hunters and professionals to monitor the movement and health of local deer populations, but estimating the age of white-tailed bucks from camera imagery remains challenging. One technique known as “Aging On The Hoof” (AOTH) attempts to determine age by analyzing the location and date of each image as well as the relative body proportions of the buck in the image [1]–[5]. When the buck’s body measurements are not known, human AOTH estimate averages just 36% – less than half the accuracy required for management-related selective harvest decisions (≥ 70%) or research purposes (≥ 80%) [6].

However, when the buck’s body measurements are known, morphometric models can be used to achieve a prediction accuracy of 63% during post-breeding periods [7]. More recent efforts have achieved significantly greater prediction accuracy (76.7%) by applying computer vision (CV) and deep learning (DL) to trail camera imagery [8], providing the first automated approach that meets the accuracy thresholds necessary for practical wildlife management applications.

### 1.2 Post-mortem dental analysis

Post-mortem dental analysis (PDA) offers another approach to aging white-tailed deer, comprising two techniques: tooth replacement and wear (TRW) and cementum annuli (CA). First introduced in 1949, TRW uses visual characteristics of the deer’s teeth (ex. tooth count, ridge structure, and material structure) to estimate the deer’s age [9]; an illustrated set of TRW criteria was provided in 1980 by Larson and Taber [10]. In practice, TRW is a two-part technique – tooth replacement enables enthusiasts to discriminate between fawns, 1.5 year old deer, and deer 2.5 years or older. The second aspect, tooth wear, enables further age separation from 2.5 years into the older age classes [11].

CA, on the other hand, counts the number of rings formed by seasonal eating patterns within thin slices of a deer’s tooth [9], [12]–[20]. Formed by cementum deposition on the roots of the teeth each year, light portions of each ring are formed in summer months and the dark portions are formed in winter months [10]. Ring patterns are not affected by the sex of the white-tailed deer, and do not change with the physiological states associated with rut or pregnancy [21].

While TRW is criticized based on the variation of tooth wear from abrasion of the deer’s food and soil types, malnutrition, and biologist’s training [14]–[16], CA has also been shown to fail due to indistinct, incomplete, or condensed annulus patterns [13], [16], resulting in inconsistent age estimates of paired incisors from the same deer [20]. The flawed nature of both methods means that CA cannot be used as the validated age of TRW estimates, and vice versa.

Nonetheless, both methods have been covered exhaustively in the literature for white-tailed deer, and some studies even report similarities in age prediction error rates between mule deer, white-tailed deer, and elk [18]. TRW and CA analyses have also been performed over many geographic regions, including the United States (Alabama [22], Mississippi [23], Montana [18], South Carolina [19], Texas [16], [24], and Virginia [15]), and Canada [14].

Despite the tendency to focus solely on male white-tailed deer [20], [24], studies have been performed on both sexes [22] showing little difference in CA age determination accuracy [18], [21]. CA measurements have been taken using the first, second, and third molars (*M*_1_, *M*_2_, and *M*_3_, respectively) [13], [14], [19], the primary incisor (*I*_1_) [14], [16], and the third premolar (*P*_3_) [24]. Additionally, researchers have noted that annulations were more apparent and consistent in the cementum than in the dentine [14].

Comparisons between the two methods have also been presented with mixed results. Some studies indicated that CA and TRW produced similar accuracies [17], while others report higher accuracy with CA [18], [24]; still others have reported TRW to be the better performer even when the CA technique is performed in a specialized laboratory [16]. Additional studies show congruence variation between TRW and CA as a function of age [15], [20], suggesting wildlife managers use a mix of the two methods depending on the age of the deer.

Recent computational approaches to deer aging have included regression analysis of cementum measurements and K-Nearest Neighbor (KNN) analysis of tooth wear patterns. While Cooper et al. found a linear relationship (*r*^2^ = 0.727) between age and dentine width [24], Meares et al. determined that dentine-to-enamel ratios (DER) could not reliably separate deer aged 2.5-4.5 years due to individual variability [19], suggesting limited improvement over visual assessment methods.

## 2 Teaching TRW

Unlike AOTH, data used in PDA are obtained only after the deer’s death, severely limiting availability. Jawbone data from past studies is rarely made available to the public, and no central repository exists for images of known-age white-tailed deer jawbones. If jawbone specimens of known-age deer become available, professional laboratories can take weeks to process a single sample [11], resulting in a sizable backlog of requests.

To enhance awareness within the outdoor community and alleviate the workload on dental analysis experts, many organizations provide tutorials on best TRW practices. In each case, the tutorial showcases jawbones from known-age white-tailed deer of different ages; for each age, the corresponding jawbone is examined and an explanation of the deer’s dental structure is provided. As more examples are presented in the literature, enthusiasts and professionals alike are able to learn and develop an intuition for predicting a deer’s age based on their own observations.

A typical white-tailed deer jawbone is illustrated in Figure 1. The annotations indicate the first, second, and third premolars (*P*_1_, *P*_2_, and *P*_3_) in yellow, and the first, second, and third molars (*M*_1_, *M*_2_, and *M*_3_) in white. Finer details and structures of individual teeth are identified in the inset, including the cusp, crest, enamel, and dentine.

**Figure 1.**
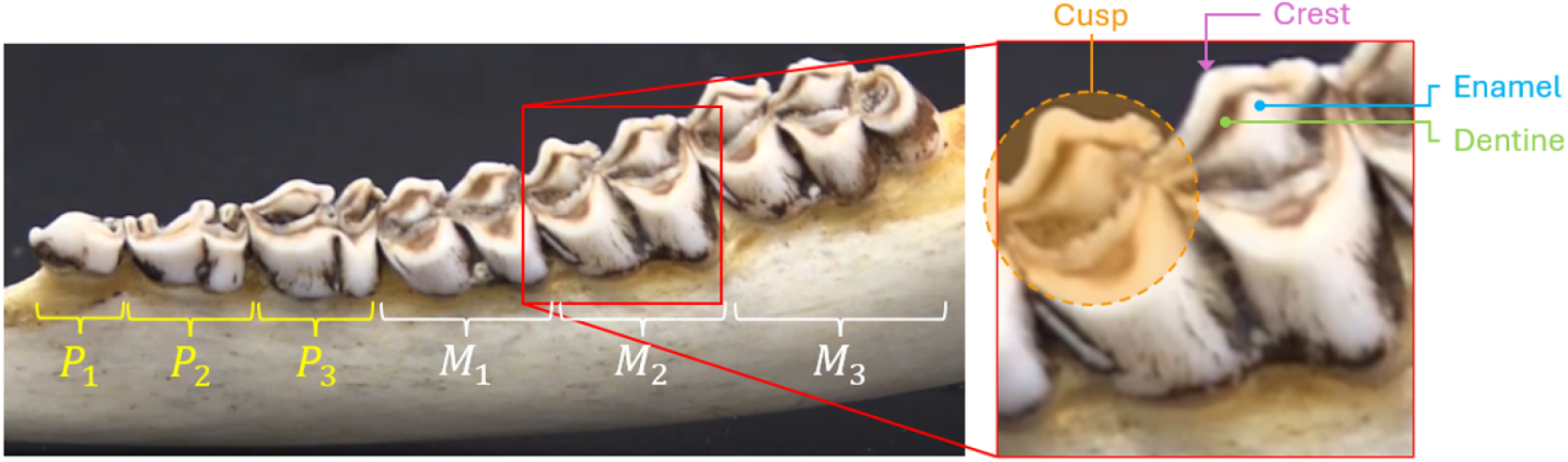
An image of a deer jawbone with an inset illustrating detailed features.

TRW analysis of each jawbone follows a sequential decision tree to determine age based on dental characteristics, including the DER of specific teeth. Analysts work through each step in order, stopping when a condition is not met. When this occurs, the deer’s predicted age corresponds to the last satisfied step. If all conditions are met, the deer is classified as 6.5 years or older. The TRW decision tree presented in many resources resembles the following:

1. Count the number of teeth in the jawbone. Newborn deer will only have four teeth.
2. If there are fewer than six teeth but more than four, the deer is a fawn (0.5 years old).
3. If there are six teeth, count the crests on *P*_3_. If three crests are present, the deer is 1.5 years old.
4. If *P*_3_ has two crests, the deer is at least 2.5 years old.
5. Examine the DER of the sharp ridges of the inside portion of *M*_1_. If 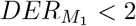, the deer is 2.5 years old.
6. If 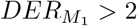 and 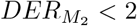, the deer is 3.5 years old.
7. If 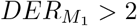 and 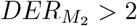 and 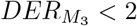 the deer is 4.5 years old.
8. If 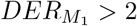 and 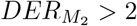 and 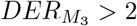, the deer is at least 5.5 years old.
9. Re-examine *M*_1_. If *M*_1_ has started to flatten, the deer is at least 6.5 years old.

It should be noted the steps listed above also contain caveats. For instance, a jawbone satisfying steps 1-4 could still represent a 1.5 year old buck if the pre-molars are clean and unstained [25]. Additionally, steps 5-9 heavily rely on the DER of the molar teeth, but this condition has been rendered questionable by Meares et al. [19].

## 3 Data

Just as computer vision models have successfully estimated the age of white-tailed bucks in trail camera imagery [8], this study uses a similar approach with jawbone analysis. By constructing a database of jawbone images from known-age deer, a deep learning model was built to predict age directly from dental imagery. The multi-fold ensemble utilizes Convolutional Neural Networks (CNN) to automatically identify and extract relevant features from these images, eliminating the need for manual tooth replacement and wear evaluation. Importantly, no TRW decision rules or dental aging guidelines were provided to the model – the CNN autonomously discovered the relevant dental features and age-related patterns through data-driven learning.

Taken from online videos, blogs, and other media, the image database compiled for this study contains 243 colored images of jawbones from white-tailed deer. Seventeen independent sources were used to collect the data, including the Quality Deer Management Association (QDMA), National Deer Association (NDA), state park websites, wildlife offices, and universities, each containing one or more highly experienced wildlife professionals. No special consideration is given to sex or geographic location, nor was any one source given superiority – each source’s images and ages are represented within the dataset.

Each image undergoes a standardized preparation process, shown in Figure 2. Once captured, image processing algorithms on a Samsung Galaxy S25 Ultra are used to remove writing or other markings that may provide extra information to the CV algorithm. Although fingertips were removed from some images, they were left in other images to mimic an enthusiast submitting images in the field. The resulting image is then cropped to a rectangular 2:1 aspect ratio, ensuring all teeth within the jawbone are included in the image. Backgrounds of the original image are left untouched to ensure the CV model learns to extract useful biological information and ignore distracting background clutter or variation.

**Figure 2.**
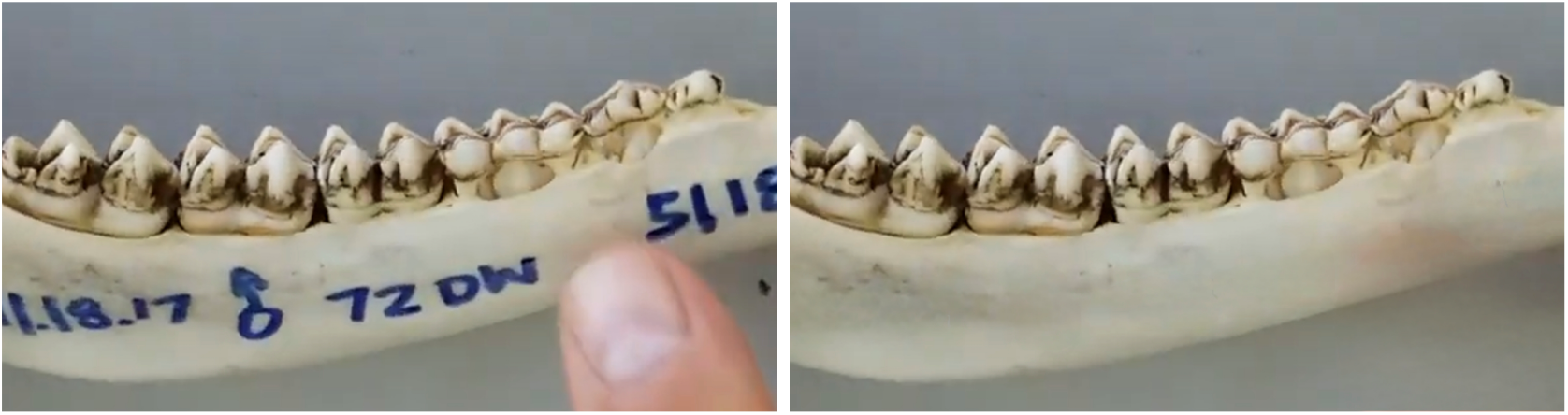
(Left) The original screen capture from a PDA tutorial is processed to produce (right) a cleaned and resized version of the original image.

A subset of processed images is shown in Figure 3, illustrating the variety of orientations, lighting conditions, and backgrounds. Since none of the images contain artificial borders or digital artifacts, any classification accuracy achieved by the model reflects its ability to recognize genuine biological age markers.

**Figure 3.**
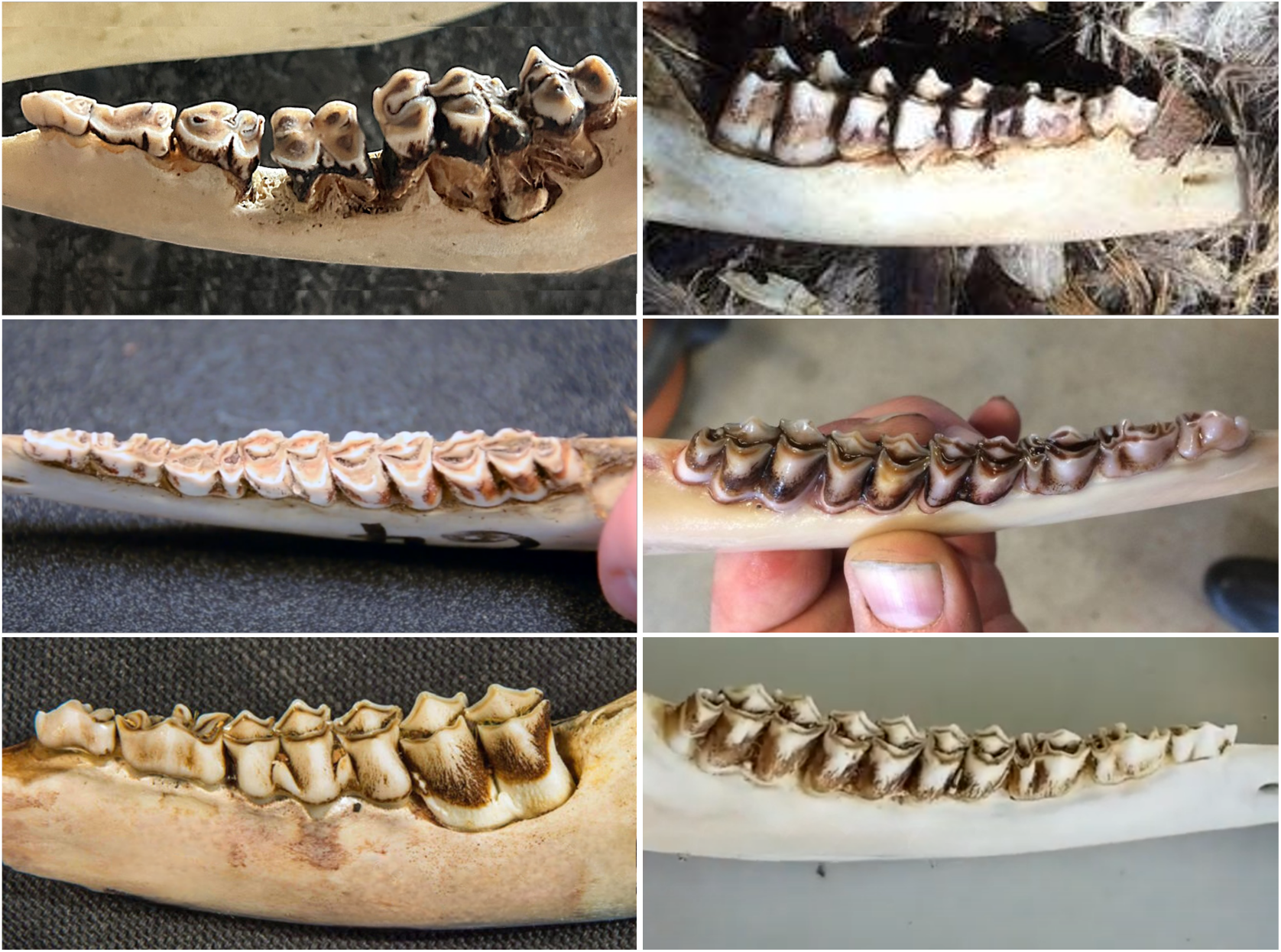
A sample of images from the standardized dataset.

### 3.1 Issues and availability

Once the data are standardized, two age-related issues remain. The first issue, lack of precision in record keeping of known-age deer, arises due to the age recording process for deer jawbones. Due to the life cycle of white-tailed deer, age is often estimated in half-year increments (ex. 0.5, 1.5, etc.). Newborn fawns are born in the Spring and mature deer are often harvested during the Fall hunting season. Even though white-tailed deer have been known to live longer than 20 years [26], deer that exceed 5.5 years of age are typically grouped together as “5.5+ years”. In a similar manner, deer within this study’s dataset that exceed 5.5 years of age are artificially labeled as 5.5 for analysis. Similarly, jawbones from newborn deer are artificially labeled as 0.5 years old.

The second challenge involves unequal representation across age classes. While PDA datasets benefit from community-wide interest in deer of all ages, age distribution remains uneven across the six age classes (0.5, 1.5, 2.5, 3.5, 4.5, and 5.5+ years). Data augmentation techniques are used to address the imbalance.

### 3.2 Age distribution

The bar chart in Figure 4 shows the age distribution in the non-augmented dataset; numbers above each bar indicate the number of samples within that age class. For example, the database contains 39 images of 0.5 year old deer, and 62 images of 1.5 year old deer. Jawbone images of each deer aged older than 9.5 years were sourced from National Deer Association media.

**Figure 4.**
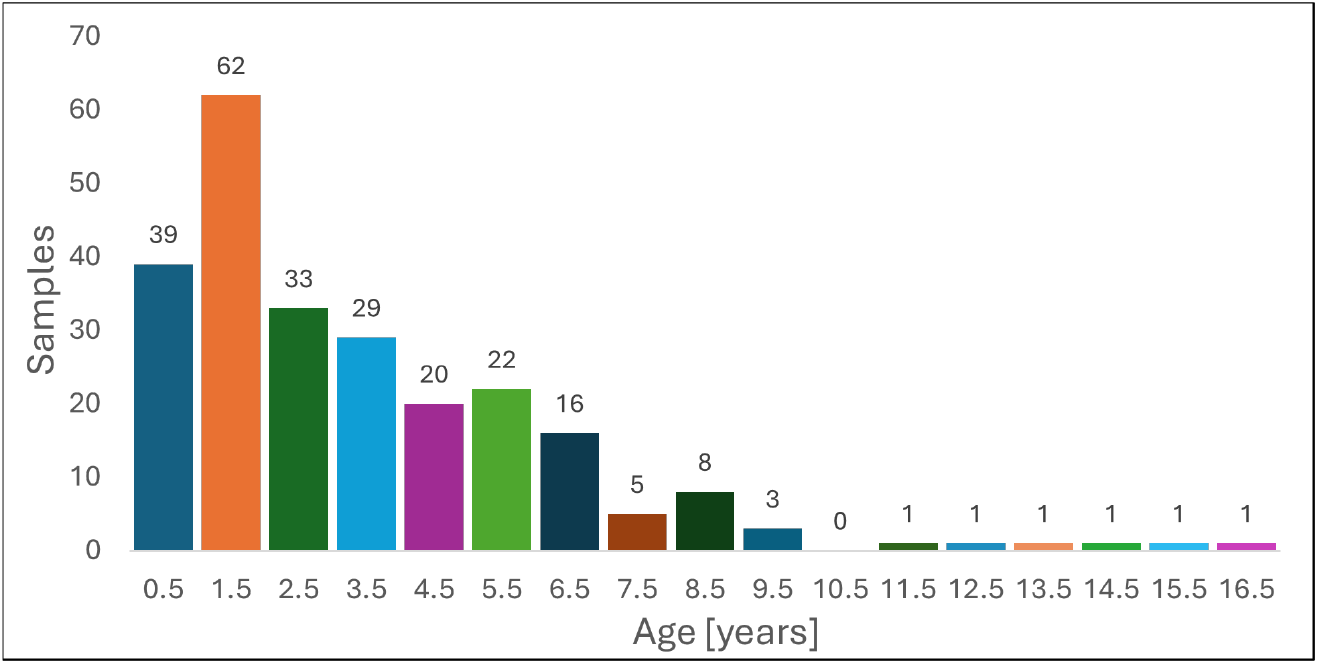
A plot of the age distribution of white-tailed deer within the dataset.

### 3.3 Augmentation

Twenty percent of the dataset was reserved as a held-out test set, with the remaining 80% comprising the training dataset. Stratified sampling ensured proportional representation of age classes across both partitions, maintaining distributional integrity for unbiased evaluation. Deep learning classification tasks typically require approximately 1,000 samples per class for optimal performance, representing a 25-fold increase over the sample sizes available in this study.

Data augmentation increases the training dataset size by applying realistic modifications that preserve jawbone characteristics and proportions. These modifications mirror natural field variations: brightness adjustments simulate different lighting conditions, rotations account for various jaw orientations during photography, and horizontal flipping represents imaging either side of the jaw. Since augmentation occurs independently for each age class, this approach enables perfect class balancing across age groups before model training.

However, excessive augmentation risks destroying age-distinguishing features. For instance, over-cropping may eliminate key tooth characteristics, while extreme brightness adjustments can obscure dental details that are critical for accurate age classification.

### 3.4 Limitations

This study experienced several limitations. First, potential specimen-level data leakage may occur due to multiple images from the same resource being split across training and test sets. While the attention map analysis (Section 5.2) suggests the model learns biologically relevant features rather than specimen-specific artifacts, the 13.1% performance gap between cross-validation and test accuracy indicates some overfitting. Future work in this field could implement specimen-level splitting to completely eliminate similar data leaks.

Second, our dataset of 243 images, while representing a substantial collection effort, is modest compared to typical deep learning applications. However, this reflects the practical reality of wildlife research data availability outside major institutional collections.

## 4 Computer Vision

### 4.1 Model architecture

Given the geometric complexity of the teeth and jawbones, transfer learning was chosen to take advantage of pre-built CNN models trained on the ImageNet database. Illustrated in Figure 5, the multi-fold ensemble approach was selected based on previous studies demonstrating white-tailed deer aging from trail camera imagery [8], where ensemble methods provided more robust and reliable predictions than single models.

**Figure 5.**
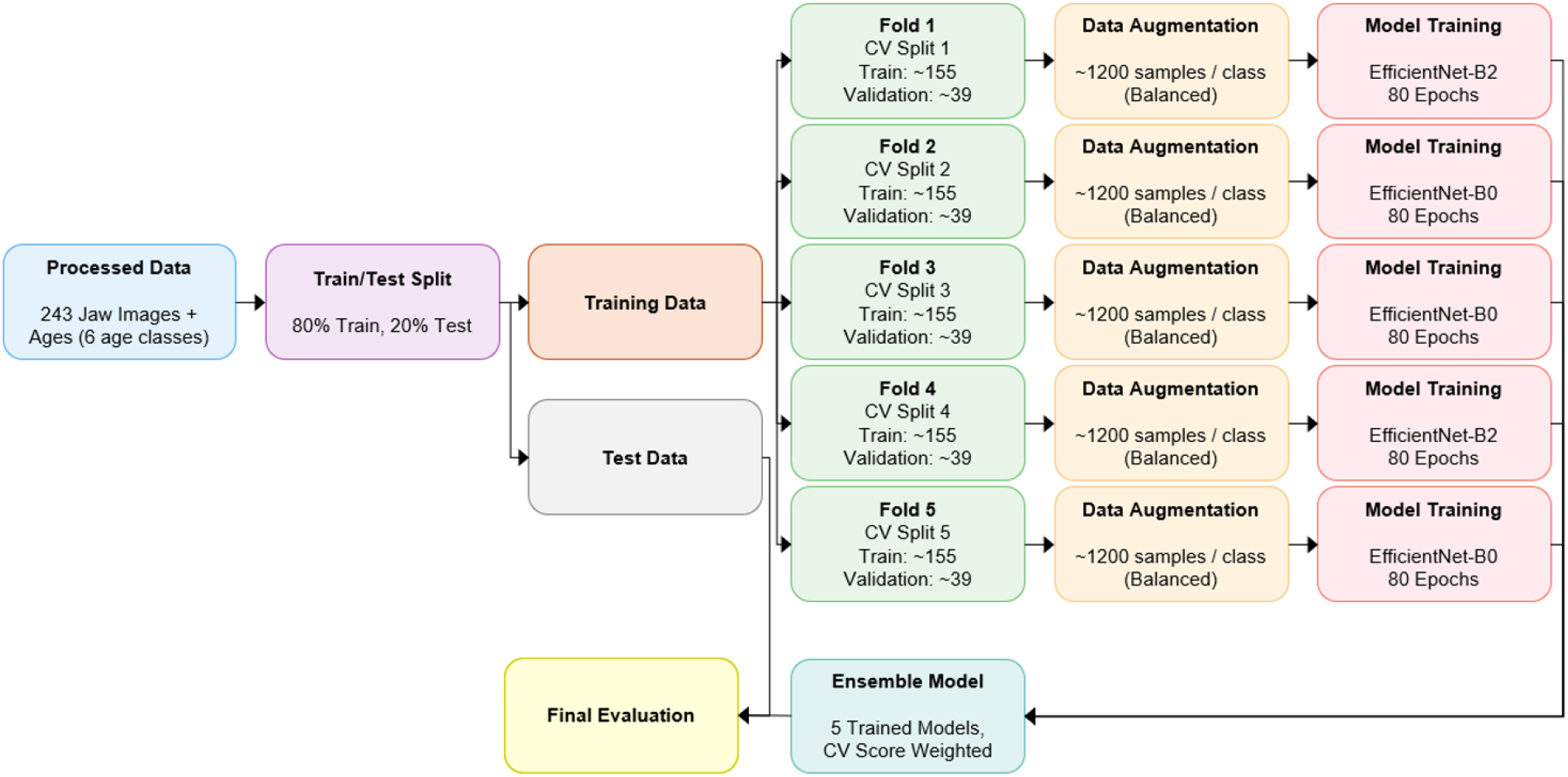
An illustration of the multi-fold ensemble training configuration.

A nested cross-validation approach was employed for model development. The training data were partitioned into five stratified folds, each preserving the original age class distribution. Each fold was subdivided into training (155 images) and validation (39 images) components. Data augmentation was then applied exclusively to the training component of each fold, expanding the sample size to 1,200 images to address class imbalance and enhance model generalization.

Each fold is subsequently sent to an EfficientNet model [27] for training; folds 2, 3, and 5 utilize EfficientNet-B0, while folds 1 and 4 utilize EfficientNet-B2. Each of the five EfficientNet models were trained using AdamW optimization with differential learning rates, 0.0003 for unfrozen backbone layers and 0.001 for the trainable classifier head. Learning rates followed cosine annealing scheduling with *T*_*max*_ = 80 and *η*_*min*_ = 1 *×* 10^*−*6^, combined with early stopping using a patience of 20 epochs. The first three blocks (blocks 0-2) of each EfficientNet architecture were frozen to preserve low-level feature representations, while deeper layers adapt to dental age classification.

Label smoothing (*α* = 0.1) and dropout (*p* = 0.3) provided regularization to prevent overfitting on the limited dataset. Cross-entropy loss served as the primary objective function, with mixed precision training accelerating convergence. Architecture selection was performed per fold, choosing the best-performing variant among EfficientNet-B0, B1, and B2 based on validation accuracy. Training typically converged after approximately 40-60 epochs per fold, with total ensemble training time of approximately 536 minutes on NVIDIA RTX 2060 hardware. Ensemble accuracy is tested on unseen data from the isolated test image set.

To arrive at a final classification prediction, the inference phase (Figure 6) uses Test-Time Augmentation (TTA) – a method that averages predictions made on the original input and its horizontally flipped version to arrive at a final prediction value, improving the ensemble’s overall robustness and accuracy.

**Figure 6.**
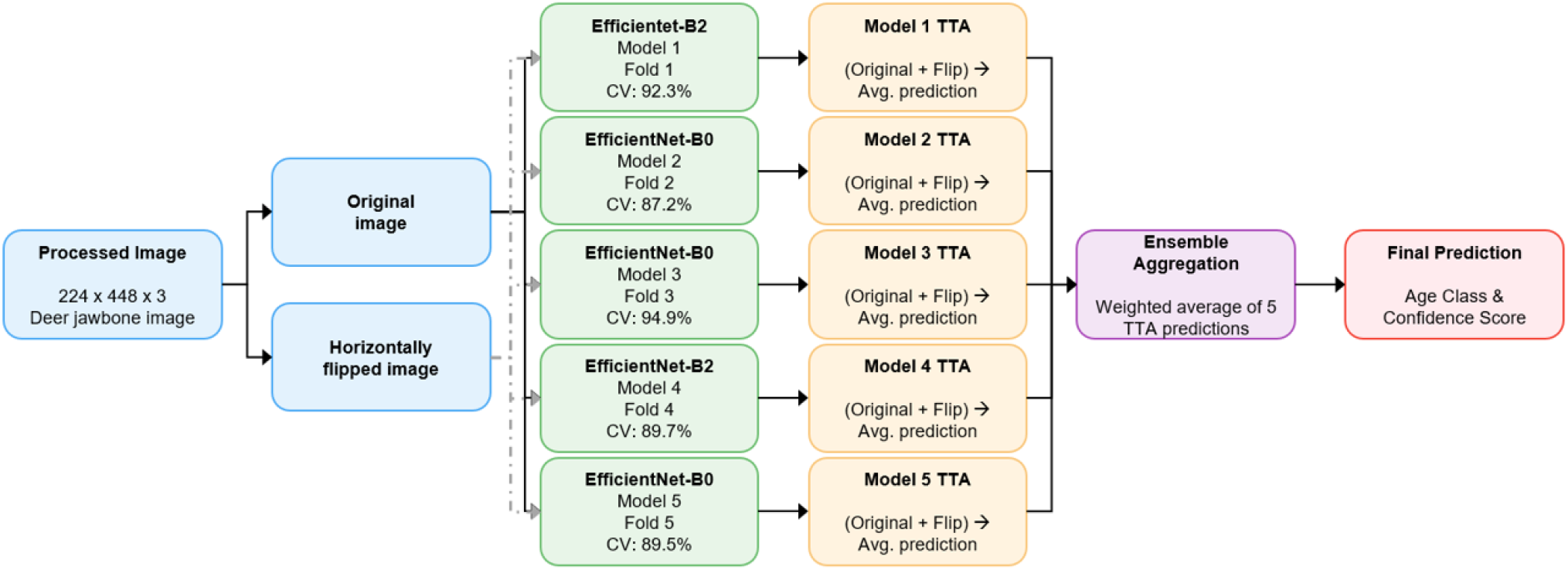
An illustration of the multi-fold ensemble model.

### 4.2 EfficientNet

The B0 and B2 versions of EfficientNet are similarly structured, but differ in their scaling parameters. As illustrated in Figure 7, EfficientNet contains seven sequential MBConv blocks (0-6) plus a head and classifier. When a colored image is fed to an EfficientNet model, the stem convolution layer spatially downsamples the input image from 224×448 pixels to 112×224 pixels, while also performing initial feature extraction, converting the three-channel RGB image into 32 feature maps that highlight basic visual patterns.

**Figure 7.**
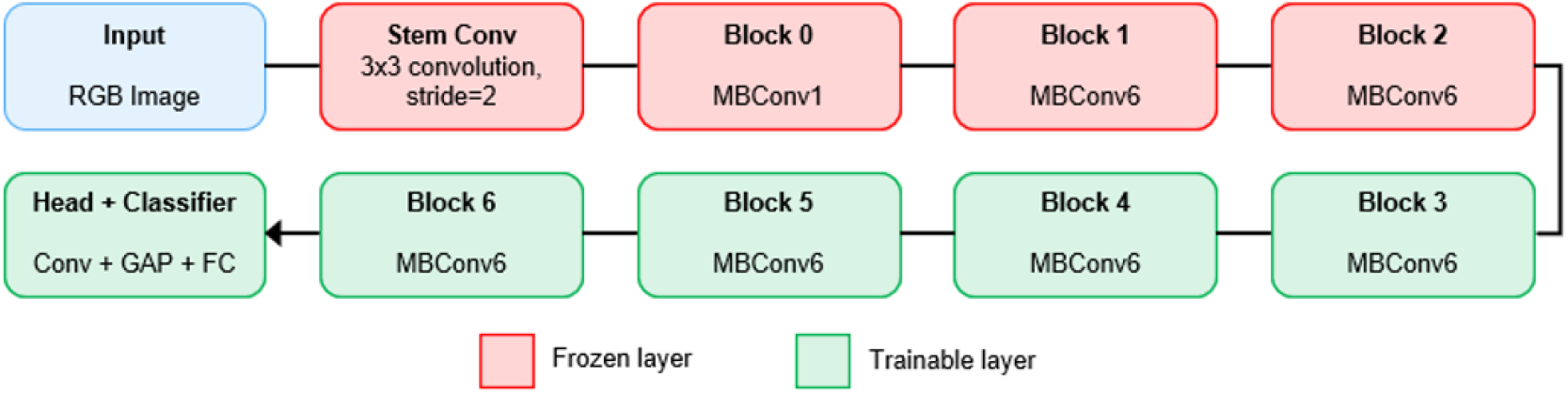
The general structure of the EfficientNet model.

These feature maps are then processed through successive MBConv (Mobile Inverted Bottleneck Convolution) blocks; the number after “MBConv” (ex. MBConv*6*) indicates the number of channels that are expanded during processing. Block 0 extracts fundamental features such as edges, lines, and simple textures. Block 1 identifies low-level patterns including corners, simple shapes, and basic texture combinations. Block 2 combines these into mid-level features that capture small object parts and more complex texture relationships. These first three blocks remain frozen during training because their generalized feature extraction capabilities transfer effectively across diverse image domains, providing a robust foundation for domain-specific learning in the subsequent trainable blocks.

Unlike blocks 0-2, blocks 3-6 remain trainable to enable domain-specific feature learning for dental age assessment. Similar to the frozen blocks, blocks 3-6 employ MBConv6 architecture but adapt their learned representations to dental-specific characteristics. Blocks 3 and 4 focus on recognizing dental-specific shapes (e.g., tooth outlines, root structures) and complex wear patterns (e.g., cusp morphology, enamel wear surfaces). Blocks 5 and 6 perform feature integration (e.g., multi-tooth eruption analysis, jaw-level wear patterns) and age-specific classification (e.g., combining dental signatures into age categories). The hierarchical learning progression through these trainable blocks can be conceptualized as: Block 3 identifies individual dental structures, Block 4 detects wear characteristics, Block 5 integrates multi-tooth patterns, and Block 6 maps these complex features to specific age classifications (e.g., 2.5 years).

Block 6 transfers 320 feature maps of complex dental age patterns to the head for feature refinement. The head applies a 1×1 convolution layer followed by Global Average Pooling, which converts the spatial feature maps (320×7×14) into a single 1280-element feature vector by averaging across spatial dimensions. The condensed feature vector is then passed to the classifier, which applies dropout regularization and a linear transformation to produce the final age classification probabilities across the six age categories. Table 1 details the fine structural differences between EfficientNet-B0 and EfficientNet-B2.

**Table 1.**
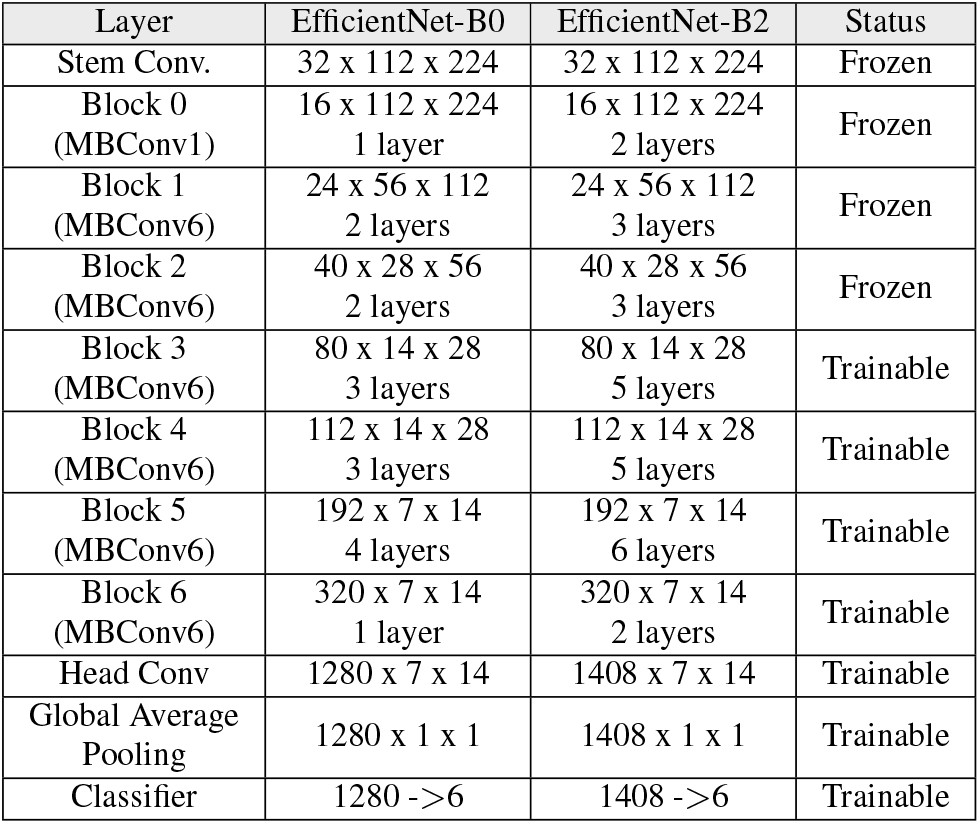
Structural details of the EfficientNet-B0 and EfficientNet-B2 models.

## 5 Results

### 5.1 Accuracy metrics

Deep learning model accuracy is often described in terms of the model’s precision, recall, and F1 score. Precision measures the reliability of the model’s predictions for a specific age class, while recall measures the model’s ability to find all deer of a specific age class. The F1 score provides a harmonic mean between precision and recall, acting as a single metric that balances both the model’s reliability (precision) and completeness (recall). The bar chart in Figure 8 shows the ensemble’s F1 score for each class, highlighting the model’s difficulty in correctly predicting ages for 2.5 and 3.5 year old deer; the dashed red line indicates the F1 score’s macro average (0.760).

**Figure 8.**
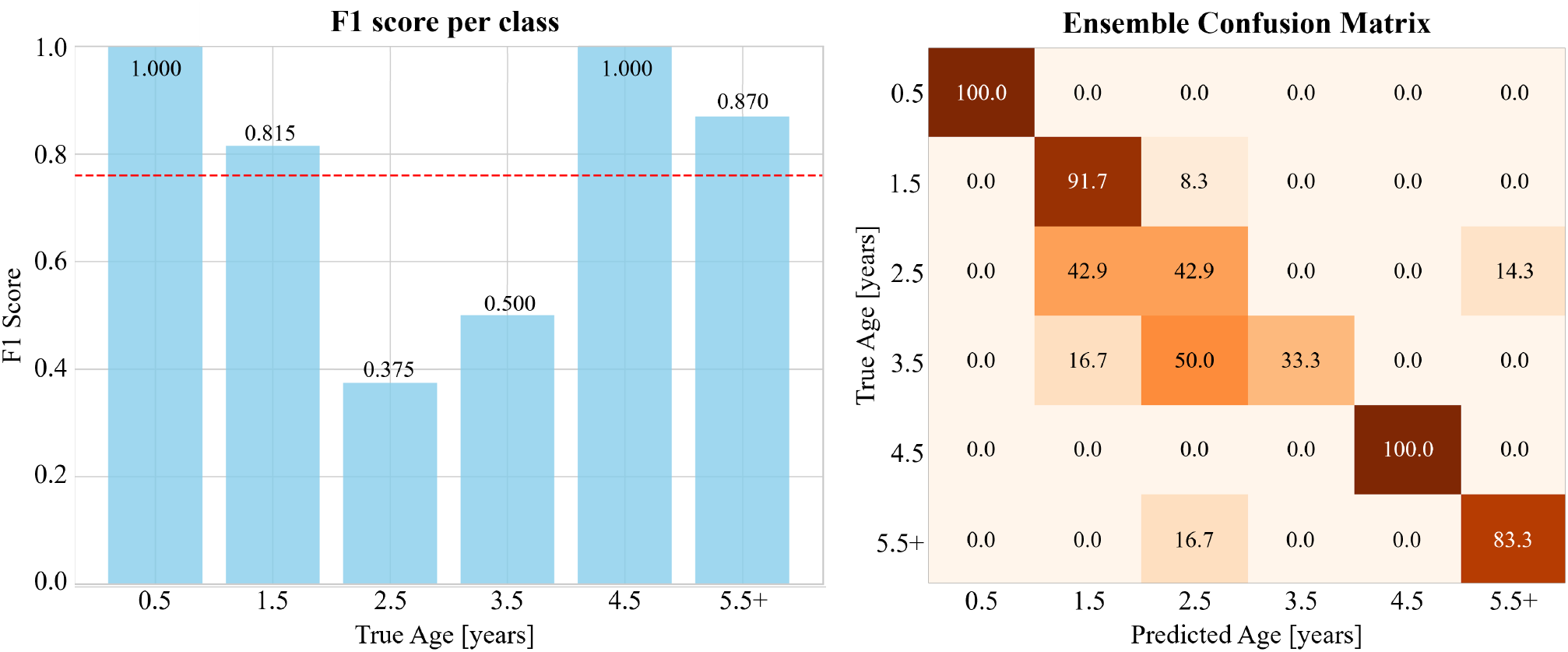
Accuracy metrics for the EfficientNet ensemble are illustrated via (left) F1 score per class and (right) a confusion matrix.

The ensemble’s confusion matrix (Figure 8 (right)) further details the ensemble’s difficulties with these age classes. For a true age of 2.5 years, the ensemble tends to guess evenly between 2.5 and 3.5 years of age. But when the true age class is 3.5 years old, the model predicts 2.5 years old half the time, only correctly guessing the true age one third of the time.

Across all age classes, the EfficientNet ensemble achieved an average cross-validation accuracy of 90.7% and a test accuracy 77.6%. The 13.1% discrepancy between the two may be due to variance within the small test set or specimen-level overfitting, even though the model’s biological feature focus indicates it primarily learns age-relevant dental characteristics.

### 5.2 Attention heatmaps

In addition to predicting the correct ages, it is equally important to demonstrate which features the EfficientNet model is picking up on within each image. Figure 9 shows one image from each age class and its associated overlaid attention map.

**Figure 9.**
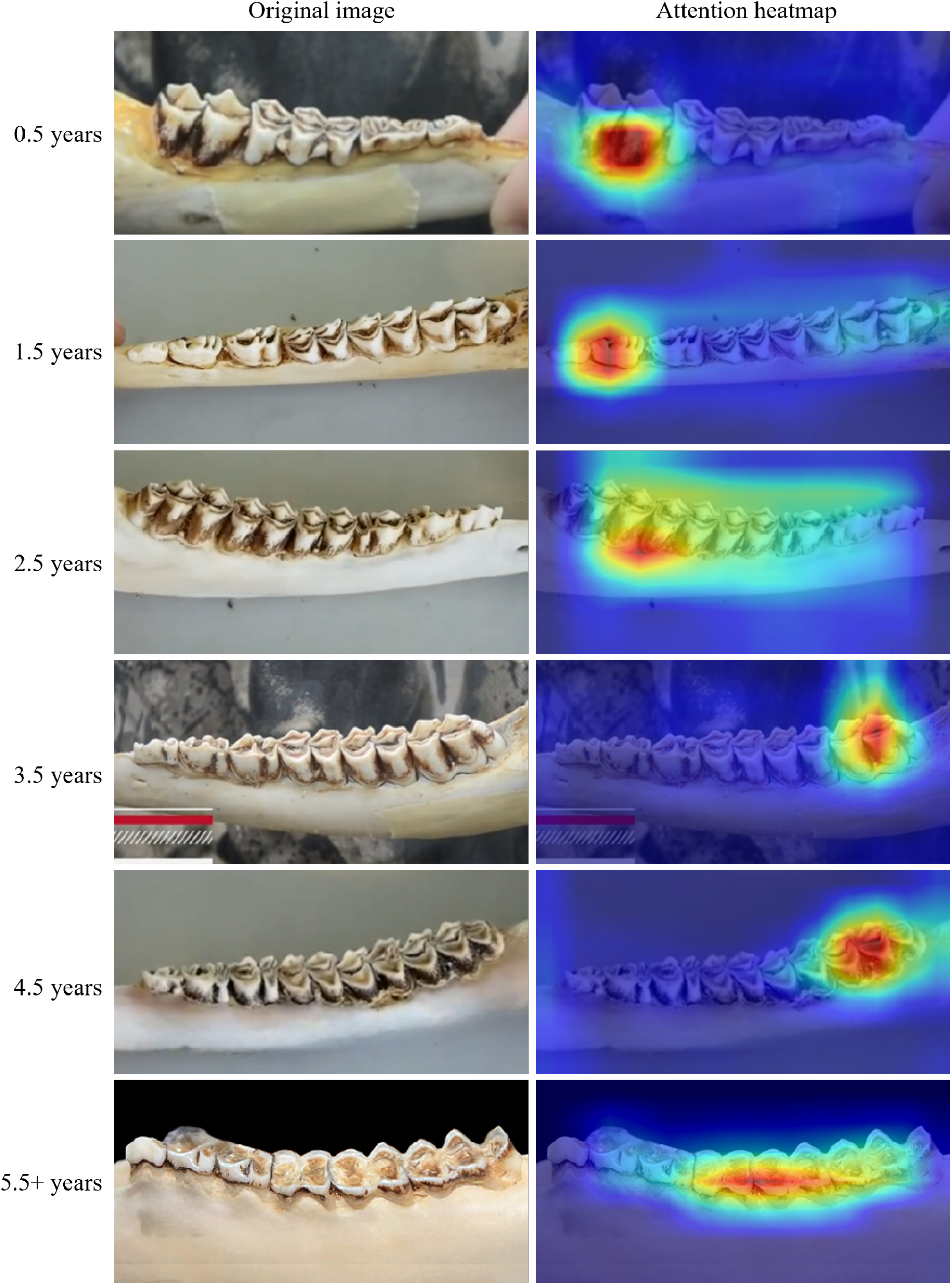
Attention map analysis showing model focus areas for each age class. The left column shows original images, the right column shows attention heatmaps overlaid on the original images. Red/yellow regions indicate high attention, blue regions indicate low attention.

Despite changes in background, finger presence, jawbone orientation, and viewing angle, the attention visualizations provide compelling evidence that the model learns biologically meaningful features. Notably, the model’s focus areas align remarkably well with the traditional TRW decision tree:

- For fawns (0.5 years), attention concentrates on the molars, consistent with tooth count assessment
- For yearlings (1.5 years), focus shifts to premolars, matching the crest-counting criteria
- For mature deer (3.5-4.5 years), attention emphasizes molar teeth, with 3.5-year focus on posterior molars and 4.5-year attention distributed across the molar region, consistent with DER evaluation patterns
- For aged deer (≥ 5.5 years), the model highlights extreme wear and flattening characteristics

This biological plausibility of learned features provides confidence that the model captures genuine aging signatures rather than spurious correlations.

## 6 Discussion

The 77.6% test accuracy achieved by the EfficientNet ensemble represents a significant advancement for automated deer aging, particularly given the constraints of working with limited data sources. This performance compares favorably to human TRW accuracy rates reported in the literature, which vary considerably based on analyst experience. The model’s ability to focus on biologically relevant dental features, as demonstrated through attention mapping, suggests potential for deployment in practical wildlife management scenarios. State wildlife agencies processing hundreds of jaw samples annually could benefit from rapid initial screening, with ambiguous cases flagged for expert review.

Furthermore, this work demonstrates the feasibility of developing specialized computer vision tools using publicly available educational resources, potentially enabling smaller organizations and citizen science projects to access automated analysis capabilities previously available only to well-funded research institutions.

## 7 Conclusions

This study presents the first known use of computer vision and deep learning in predicting the age of white-tailed deer based on jawbone images. Transfer learning with EfficientNet CNNs achieved 90.7% mean cross-validation accuracy and a test accuracy of 77.6%, representing significant improvements over traditional TRW methods and CA techniques in terms of accuracy, time efficiency, and repeatability. The technique complements previously developed AOTH CV models, providing wildlife management and outdoor enthusiasts another tool to quickly, easily, and accurately obtain white-tailed deer age estimates.

### 7.1 Future directions

Several avenues could improve upon this work including collaboration with institutional partners to access larger, specimen-tracked datasets, geographic validation using jawbones from different regions, integration with existing wildlife management databases, development of uncertainty quantification to flag ambiguous cases for expert review, and extension to other cervid species where similar dental aging techniques apply.

## ACKNOWLEDGMENTS

The author acknowledges the wildlife management organizations, state agencies, and educational institutions that provided post-mortem dental analysis training materials used in dataset construction. Particular appreciation is extended to the wildlife professionals who developed these educational resources, enabling this interdisciplinary application of computer vision to wildlife biology. Appreciation is also extended to the open-source community for the deep learning frameworks and tools that enabled this work. This research received no external funding.

## Notes

### Competing Interest Statement

The authors have declared no competing interest.

